# Single Amino Acid Change Mutation in the Hydrophobic Core of the N-terminal Domain of P22 TSP affects the Proteins Stability

**DOI:** 10.1101/2021.12.16.472976

**Authors:** Joseph A. Ayariga, Robert Villafane

## Abstract

The emergence of the severe acute respiratory syndrome coronavirus 2 (SARS-CoV-2) has significantly shifted the attention of researchers to critically investigate most viruses to understand specific characteristics that impart their virulence. For instance, the SARS-CoV-2 has undergone several mutations, with some variants classified as “variants of concern”, e.g., the Omicron and Delta variant of SARS-CoV-2 are known for their rapid transmission and antigenicity due to mutation in the Spike protein. P22 bacteriophage is a bacterial virus that has a tailspike protein (TSP) that performs similar functions as the Spike protein of SARS-COV-2. We previously carried out a site-directed mutagenesis of the P22 TSP to bear disruptive mutations in the hydrophobic core of the N-terminal Domain (NTD), then partially characterized the properties of the mutant TSPs. In this process, the valine patch (triple valine residues that formed a hydrophobic core) was replaced with charged amino acids (Asp or lysine) or hydrophobic amino acids (Leucine or isoleucine). Some of the mutant TSPs characterized showed significant differences in migration in both native and SDS-PAGE. Mutants with such disruptive mutation are known to show non-native properties, and as expected, most of these mutants obtained showed significantly different properties from the WT P22 TSP. In this work, we further characterized these mutant species by computational and in vitro assays to demonstrate the validity of our previous inference that the valine patch is a critical player in the stability of the N-terminal domain of the P22 TSP.

## 1.0 Introduction

The emergence of the severe acute respiratory syndrome coronavirus 2 (SARS-CoV-2) in 2019 has significantly turned the attention of researchers to deeply investigate most viruses to understand specific characteristics that impart their virulence **(1)**. Frequent gene mutations are typical in both animal and bacterial viruses **(2, 3)**, while some mutations are either deleterious and hence of poor evolutionary significance to the virus, some other mutations affect the functional characteristics, thus changing the virulence, disease severity or interactions with host immunity **(4)**. For instance, the SARS-CoV-2 has undergone several mutations, with some variants classified as “variants of concern,” e.g., the Omicron and Delta variant of SARS-CoV-2 are known for their rapid transmission and antigenicity **(5, 6)** due to mutation in the Spike protein. In our system, the P22 bacteriophage is a well-studied phage and has been used model phage for genetic, molecular, and structural studies **(7, 8, 9, 10, 11)**. The phage possesses tail machinery, and attached to this tail machinery is an endorhamnosidase trimeric tailspike protein (TSP) that plays the role of a receptor-finding, receptor-binding, and a receptor processing protein for the phage **(7, 10)**. It finds and binds to its host via the TSP. Interestingly the tailspike interactions with lipopolysaccharide (LPS) alone have been demonstrated to effect DNA ejection from the phage particles *in vitro* **(7)**. Published research surrounding the utility of this highly studied protein for clinical application and bacterial control is available **(12, 13)**. This endorhamnosidase can specifically degrade the lipopolysaccharides of the host bacterial envelope leading to adsorbing, invading, replication in the host, and finally lysis of the host **(14)**. The homotrimeric P22 TSP consists of three chains in which the N-terminal domain (NTD) which binds to the rest of the phage structure in the formation of a complete and infective phage **(9)**, contains 108 amino acids **(10)**. The NTD forms a trimeric dome-like structure that non-covalently attaches to the rest of the phage. While an initial study of this dome-like structure has been carried out in our laboratory, especially the influence of the first 23 amino acid of the NTD on the stability of the dome structure **(10)**, no in-depth knowledge exist on the molecular interactions occurring between this first 23 amino acids and the entire dome, and the role this 23-amino acid fragment plays in the stability of the entire globular protein. In our previous communication, we demonstrated that the hydrophobic valine patch (V8-V9-V10) located within the dome structure is critical for the stability of the NTD. We proved via site-directed mutagenesis that the valine patch and the adjacent polypeptide are crucial for interchain interaction which is necessary for the stability of the P22 TSP NTD. To further elucidate this assertion, we carried out both *in vitro* and *in silico* analysis of the variants produced by the site directed mutagenesis at the valine patch to test further and validate our previous conclusions that the valine patch is a crucial player in the stability of the NTD of P22 TSP (**10**). Both *in silico* and *in vitro* analysis confirmed our previous findings that the valine patch is crucial in the structural integrity of the NTD of P22 TSP. Such knowledge could prove critical for bioengineering purposes in which P22 TSP can be manipulated to achieve tunable affinity for the rest of the phage or chimeric protein that can be used for biosensing, pathogen immobilization, or streptavidin-coated magnetic beads for the purpose of ELISA.

## 2.0 Methods

### 2.1 In silico analysis of variant TSPs

To carry out the *in silico* analysis of the variant TSPs, the following computational methods were employed; wildtype P22 TSP’s N-terminal domain (NTD) template selection and retrieval from the RCSB database (PDB ID: 1LKT), amino acid changes to at the desired positions to form variant TSPs, modeling of variant TSPs, and structural validation and evaluation of models.

### 2.2 P22 TSP’s NTD selection

The crystal structure of the P22 phage tailspike protein’s (TSP’s) head-binding domain (PDB ID: 1LKT) has been solved at 2.3 Å **(15)**. Since our interest was to understand the effect of a mutation in this head binding domain (also called NTD) of the protein; that is, the effect of amino acid substitution in the region with the triple valine residues (termed “valine patch”) in the NTD, therefore, only the NTD of the P22 tailspike protein was selected. The crystal structure of the wildtype P22 TSP NTD (PBD ID: 1LKT), which is a dome-like structure, was retrieved from the RCSB database (https://www.rcsb.org/).

### 2.3 Amino acid changes and building of variant TSPs models

The crystal structure of the template in a PDB format was uploaded onto a modeling program (Biovia Discovery Studio, version 2020), the specific amino acids were substituted using this program. Then the sequence of the variant/mutant TSPs resubmitted to the Swiss-modeling program **(16)** to build the mutant models using the PDB file 1LKT as a template and models visualized using the Swiss PDB Viewer v 4.0.1 software **(16)**. Mutagenesis was carried out by replacing valine 8, 9, or 10 with a single aspartate, leucine, isoleucine, or lysine. The changes made are shown in Table 1 below.

**Table 1.**
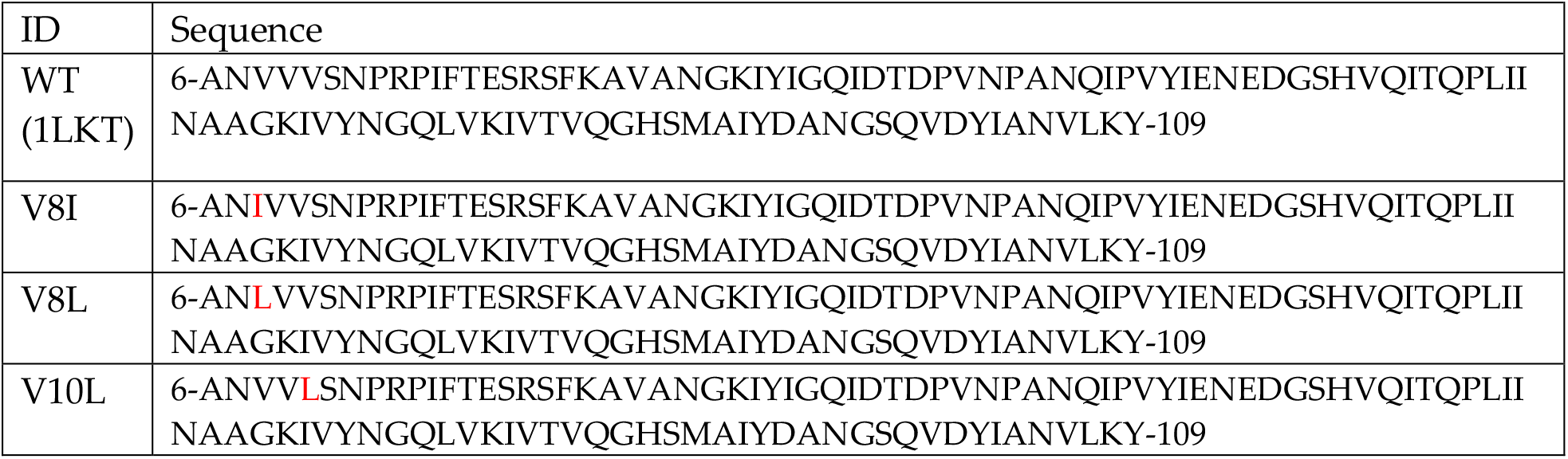
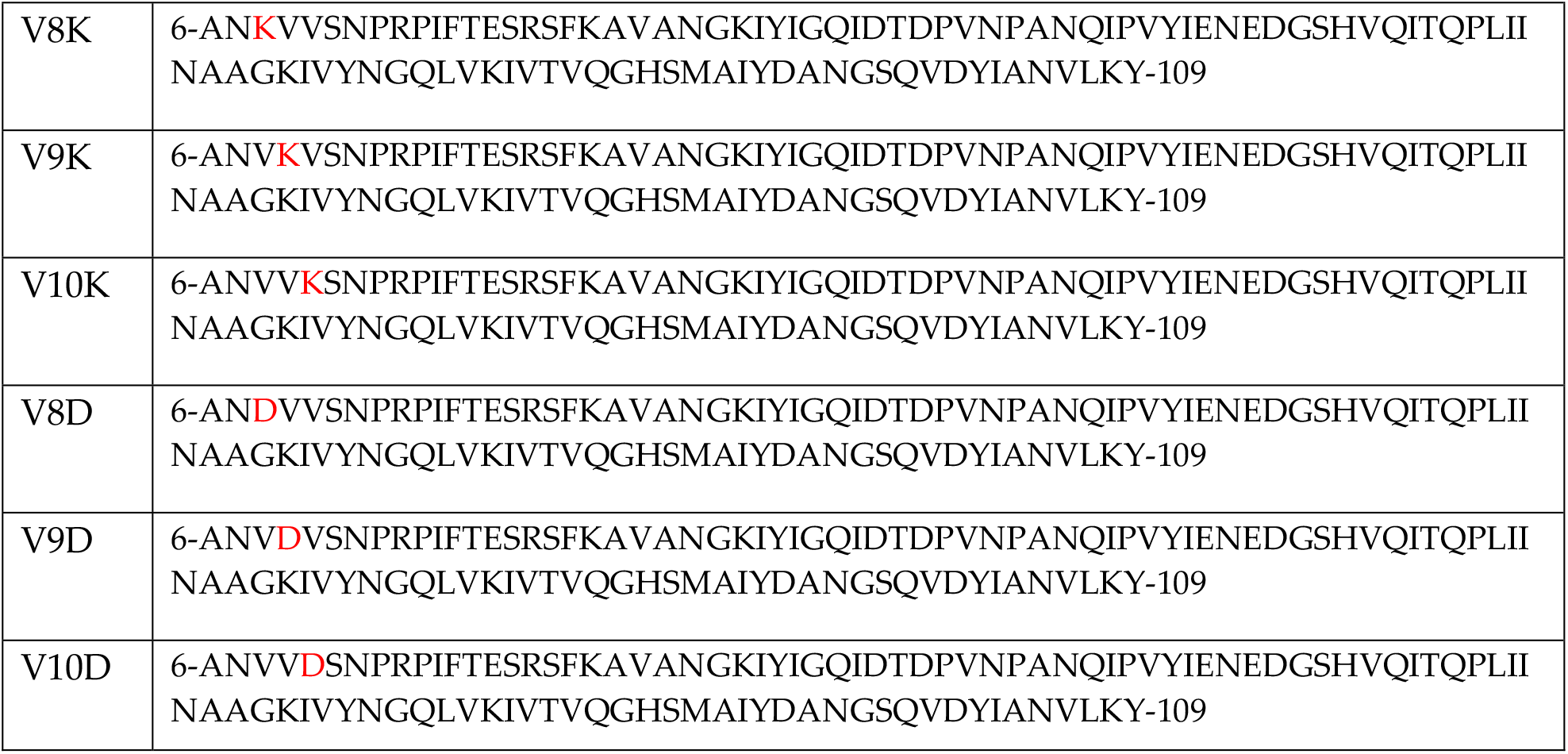
The sequences of the WT (PDB ID: 1LKT) **(15)** and variant TSPs of P22 TSP’s HBD. The mutated amino acids are shown in red. BLASTP **(17)** was used to identify homologs in the RCSB Protein Databank (https://www.rcsb.org/).

### 2.4 Structure assessment of variant TSPs using ERRAT

The 3D models of the WT and the variant TSPs created using the Swiss model tool were saved in a pdb format and submitted to SAVES v6.0 (UCLA-DOE LAB, https://saves.mbi.ucla.edu/) for the structural assessment using the ERRAT 2 program **(18)**.

### 2.5 Structure assessment of variant TSPs using ERRAT

ERRAT is a program that assesses the compatibility of protein structures by classifying atoms into three types (C, N, and O) and determines if the distribution of non-bonded interactions between atoms in a given structure matches the distribution established from a database of reliable, high-resolution structures. Therefore, the ERRAT program is an effective tool for identifying erroneous regions of a model structure. Error-values are plotted as a function of the position of a sliding 9-residue window. The error function is a derivative of the statistics of non-bonded atom-atom interactions in the given structure compared to a database of reliable high-resolution structures **(18)**. Regions of candidate protein structures that are miss-traced or misregistered can then be identified by analyzing the pattern of non-bonded interactions from each window. In this work, ERRAT2 was used to verify the variant TSPs in order to determine the level of error as well as the errenous regions of the variant TSPs generated due to the change of the WT amino acid to the variant.

### 2.6 Molecular alignment and morphing Analysis of variant TSPs

To investigate any differences in molecular motion between variant TSPs and the WT TSP, we subjected the modeled variants to molecular alignment and morphing analysis using the crystal structure of the WT TSP NTD (PBD ID: 1LKT) as a reference structure. Molecular alignment and morphing were carried out using PyMol 2 (Schrödinger, INC, https://www.schrodinger.com/products/pymol) with the start conformation being the WT NTD TSP, and the mutant models serving as the end conformation. Conformational morphing was run at refinement cycles of 10, and the outputs were set at 50. The Rigimol interpolation was adopted for the morphing process. To to determine the root mean square distances (RMSD) of the mutant models, the mutant models were aligned to the WT NTD TSP Crystal structure using PyMol 2. Many-to-one alignments were carried out with the Mutant models serving as the mobile object, whereas the WT NTD TSP served as the Target selection object. The number of cycles or the maximum number of outlier rejection cycles was set at 5, with an outlier rejection cutoff in RMS of 2.0..

### 2.7 Site-Directed Mutagenesis

Site-directed mutagenesis, using the QuikChange Mutagenesis Kit was employed for the mutagenesis of the tailspike protein, and the mutation and cloning process, as well as validation and initial characterization of the cloned variant TSPs, have been carried out and published in previous work in our laboratory **(10)**.

### 2.8 Electrophoretic analysis of variant TSPs in native PAGE

To investigate the migration of the variant TSPs in a native electrophoretic condition, aliquotes of protein samples were mixed with loading buffer consisting of 50 mM Tris-HCl, 25 % glycerol, and 0.01 % bromophenol blue. The mixed aliquots were then loaded into a 10 % polyacrylamide electrophoresis (PAGE) gel well and run at 100 volts in an SDS-free running buffer. Gel bands were visualized by Coomassie blue staining. Gel images were acquired using a Bio-Rad ChemiDoc XRS imaging system. Densitometric values were acquired using the Quantity One software, and values were expressed as percentages and plotted into graphs. Experiments were run in triplicates.

### 2.9 Treatment of variant TSPs with SDS and high temperatures

Although the P22 TSP is resistant to SDS denaturation, high temperatures, and proteases, it has been evidently demonstrated that a combination treatment of the protein with SDS and heat denature it **(10, 19, 20)**. In this study, we subjected the variant TSPs to SDS-PAGE analysis at room temperature and the combination of SDS and high temperature of 70 °C. In brief, the various variant P22 TSP were incubated for set time points in 2% SDS at pH 7.4 in 70 °C. Aliquots of treated samples were taken and mixed with a loading buffer consisting of 50 mM Tris-HCl, 25 % glycerol, 0.01 % bromophenol blue, and 2% SDS. Samples were then loaded into a 10 % polyacrylamide electrophoresis (PAGE) gel and ran at 100 volts. Gels were stained using Coomassie blue stain and imaged using Bio-Rad ChemiDoc. In a separate experiment, protein samples were incubated in 90 °C heat treatment only, or 70 °C heat treatment only to assess the mutant TSPs’ thermostability compared to the wildtype P22 TSP. Experiments were run in triplicates. Densitometric values were acquired using the Quantity one software, and values were expressed as percentages and plotted into graphs.

### 2.10 Statistical analyses

For all statistical data, values were derived from multiple measurements (from replicates of 3 experiments) and averaged; the standard deviations were evaluated using P-values of Student’s t-test (one-tailed, two samples of unequal variance, significance level α = 0.05 was used unless otherwise stated).

## 3.0 Results and Discussion

Published works have demonstrated that single amino acid mutations can potentiates major thermal instability proteins due to distortion of the hydrophobic core (**21**). To assess the effect of amino acid substitution in the hydrophobic valine patch and also on the entire NTD structure of the P22 TSP, several *in silico* and *in vitro* assays were conducted.

### 3.1 Structure assessment of variant TSPs using ERRAT

As shown in Figure 1, ERRAT plots of all the variant TSPs are shown. Regions of the structure that can be rejected at the 95% confidence level are yellow; whereas, regions of the protein structures that can be rejected at the 99% level are shown in red. As shown in Figure 1, V8K (Figure 1A), V9K (Figure 1B), V10D (Figure 1F), and V8I (Figure 1H) showed the highest erroneous regions, thus indicating that the highest structural misfits were introduced by V8K, V9K, and V10D substitutions.

**Figure 1.**
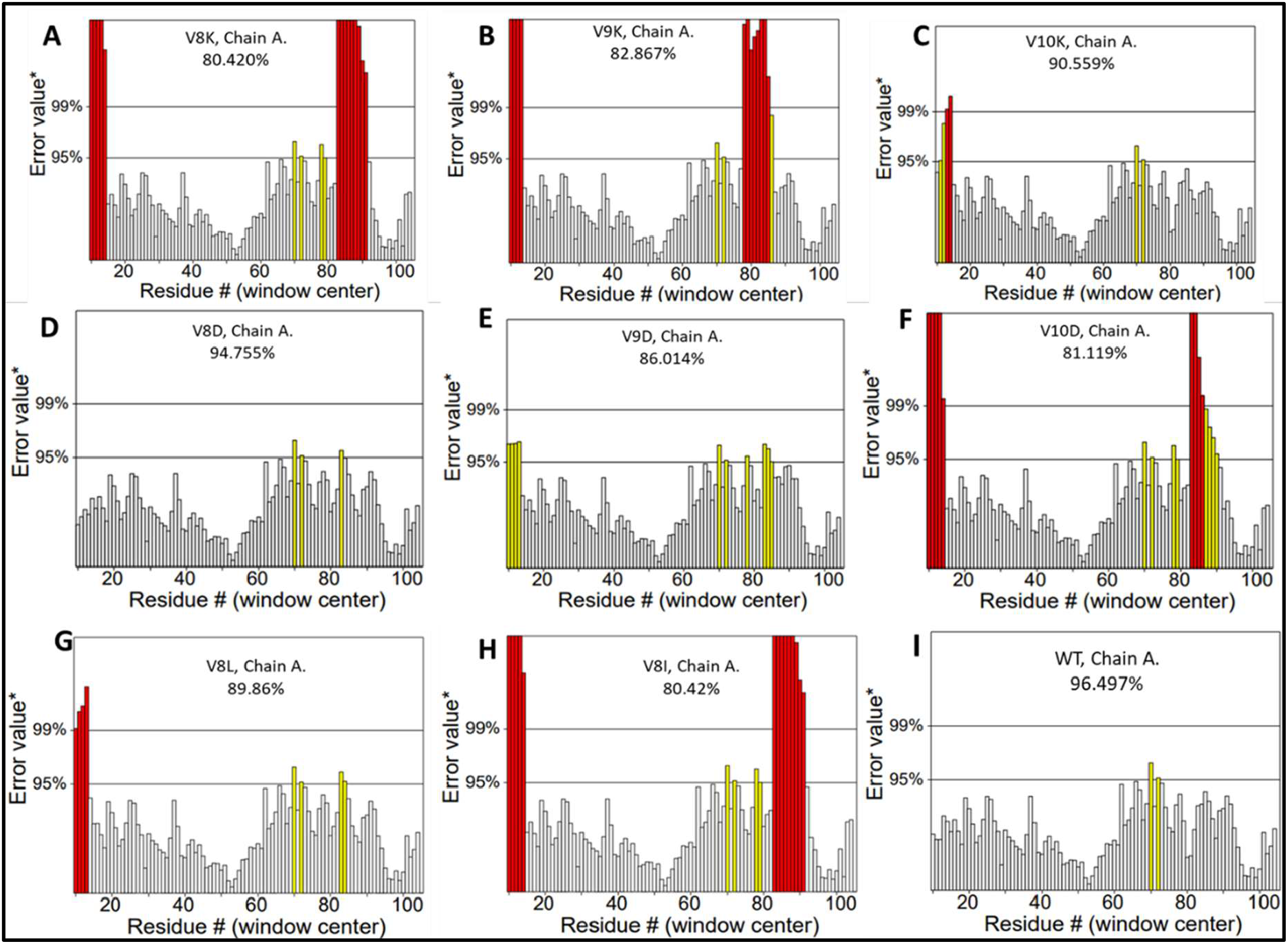
Errat plot for the variant TSP models and the WT TSP of P22 TSP. Red and yellow bars depict erroneous or misfolded regions introduced by the change in amino acid. The white bars depict regions with a lower error rate for protein structure. Regions of the structure that can be rejected at the 95% confidence level are yellow; whereas, regions of the protein structures that can be rejected at the 99% level are shown in red. *On the error axis, two lines are drawn to indicate the confidence with which it is possible to reject regions that exceed that error value. Expressed as the percentage of the protein for which the calculated error value falls below the 95% rejection limit.

### 3.2 Structure assessment of variant TSPs using Cβ Deviation

One of the crucial components for protein structure evaluation is the consideration of the geometric relationship between the Cβ and the Cα. This is due to the fact it gives information concerning the relationships between the sidechains and backbone, thus a crucial indicator of structure compatibility. For instance, if the backbone or the sidechain rotamer happens to be incompatible (misfit), model refinement compromises by distorting Cα geometry. Hence, distortion around the Cα is generally used as a sensitive method to identify misfits in protein structures. This distortion is a measure of Cβ deviation which is the ideal-geometry of Cβ from the backbone atoms, or the distance of the observed Cβ from the ideal one **(22)**. The amino acid substitution was used in creating mutant TSPs; thus, the substituted amino acid affects previous bonding and geometric space in the protein. Using Swiss modeler, an online program, we assessed the Cβ deviation for each variant created (Table 2). As shown, the highest deviation was 9, recorded by V8D and V10D. The WT TSP and V10L variant recorded the lowest Cβ deviations of 4, thus except for V10L, all variants recorded higher Cβ deviations than the WT TSP (Table 2).

**Table 2.** Scores for variant TSP models as compared to the WT TSP.

| | MolProbity | C-Beta Deviations | Bad Angles | QMEAN | Cβ | Solvation | Torsion |
| --- | --- | --- | --- | --- | --- | --- | --- |
| WT | 0.77 | 4 | 8 of 3301 | 1.09 | -1.5 | -0.44 | 1.68 |
| V8D | 0.79 | 9 | 12 of 3304 | 0.95 | -1.57 | -0.39 | 1.51 |
| V9D | 0.85 | 6 | 10 of 3304 | 1.18 | -1.37 | -0.47 | 1.74 |
| V10D | 0.85 | 9 | 16 of 3304 | 0.91 | -1.61 | -0.46 | 1.51 |
| V8I | 0.79 | 8 | 9 of 3309 | 0.94 | -1.32 | -0.43 | 1.47 |
| V8L | 0.79 | 8 | 6 of 3309 | 0.92 | -1.27 | -0.4 | 1.42 |
| V10L | 0.85 | 4 | 8 of 3304 | 1.1 | -1.25 | -0.4 | 1.6 |
| V8K | 0.85 | 8 | 10 of 3304 | 0.93 | -1.41 | -0.37 | 1.45 |
| V9K | 0.85 | 8 | 9 of 3304 | 1.25 | -1.18 | -0.47 | 1.78 |
| V10K | 1.12 | 7 | 10 of 3304 | 1.01 | -1.62 | -0.67 | 1.7 |

### 3.3 Solvation potential and Torsion of variant TSP models

The solvation energies determination using the Swiss modeler program help assess the burial status of the residues of the protein **(23)** and solvent accessibilities, which are crucial determinants in protein stability and ability to resist denaturation. The highest predicted solvation energy release was recorded on V10K (−0.67), this is followed by V9K and V9D, each recording -0.47 each, the two substitutions that produced comparable solvation energies as the native structure were both from the hydrophobic amino acids substituted, V8I and V8L, which showed -0.4 each. Unexpectedly, V8D produced the least solvation energy release of 0.39. The torsional angle potential assessment shows the possibilities of the amino acid substitution altering the native structure’s local geometry. As shown in Table 2, the highest torsion was recorded for V9K (1.78), then followed by V10D (1.74) and V10K (1.70). The two hydrophobic amino acids substitutions recorded the lowest torsion of 1.47 and 1.42 for V8I and V8L, respectively.

### 3.4 Bad angles introduced and the Cβ interaction energies of variant TSP models

As shown in Table 2, the highest number of bad angles recorded was 16 out of 3304 bonds for variant TSP V9D; the lowest number was recorded for the hydrophobic amino acid substitutes V8I (9 out of 3309), V9L (6 out of 3309). The Z-scores of Cβ interaction energies of the variant TSPs demonstrated that V10K and V10D recorded the highest energies of - 1.62 and -1.61, respectively. The lowest values were recorded for the two hydrophobic amino acids substituted variants, is V8I, V8L, and V10L, demonstrating Cβ interaction energies of -1.32, -1.27 and -125, respectively.

### 3.5 Molecular morphing of variant TSPs

Molecular movement plays a vital role in protein structure and biological functions **(24)**. To gain insight into the possibility of different molecular motions that can be imparted by the amino acid changes made in the variant models, the variant models were subjected to molecular morphing using PyMol 2, with the WT TSP NTD crystal structure (PDB ID: 1LKT) serving as the reference structure. As shown in Supplementary Figures S1 to S6, it is evident that the amino acid substitution in the valine patch, which is a hydrophobic region, significantly produced varied motions. While V8D (Figure S1 and S2) produced a very small motion in the local region of the substitution, there was an observed significant increase in motion in V9D (Figure S3) and V10D variant models (Figure S4 and S5). The most dramatic and significant movement was recorded for V8K variant (Figure S6). However, the trajectories of V9K (Figure S7) and V10K (Figure S8) unexpectedly showed very small movements. The hydrophobic amino acids substitutions provided characteristically varied motions, whereas V8I variant model (Figure S9) produced a very large-scale molecular movement in the local geometry, the V8L variant model (Figure S10) produced small movement in the local geometric space. These differences might be attributable to the position of the R group in the amino backbone. Leucine which is 2-amino-4-methylpentanoic acid, has the R group positioned on carbon 4, whereas the isoleucine, which is 2-amino-3-methylpentanoic acid, has the R group positioned at the third carbon. It is quite interesting how this minuscule change could produce such a significant difference in the molecular dynamics of the local as well as the global structure of the protein. The large-scale conformational changes observed in the charged amino acids substitution mutants such as the Valine – to - K or the Valine-to-D might be due to the introduction of charged/polar groups into a purely hydrophobic region. To address this further, we measured the RMSD of these variant models after structure alignment (Table 3).

**Table 3.**
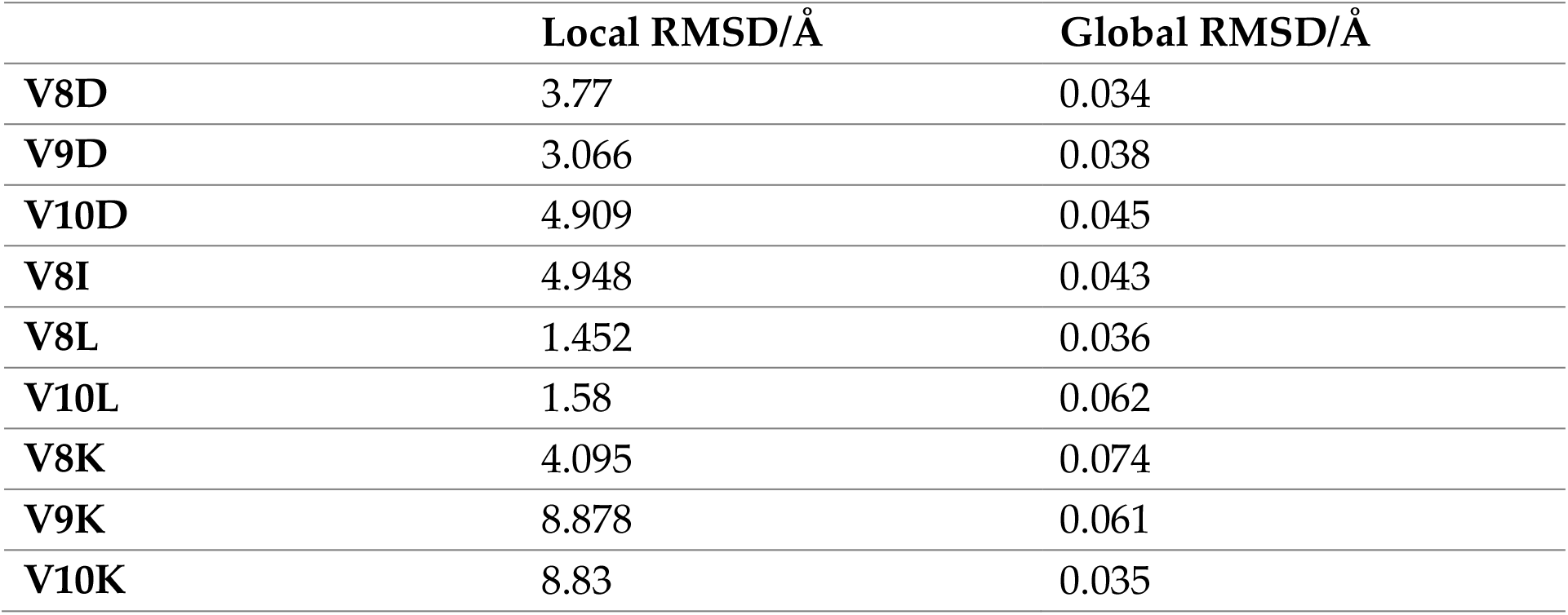
Global and local RMSDs of variant TSPs measured after amino acid substitution.

|  | Local RMSD/Å | Global RMSD/Å |
| --- | --- | --- |
| **V8D** | 3.77 | 0.034 |
| **V9D** | 3.066 | 0.038 |
| **V10D** | 4.909 | 0.045 |
| **V8I** | 4.948 | 0.043 |
| **V8L** | 1.452 | 0.036 |
| **V10L** | 1.58 | 0.062 |
| **V8K** | 4.095 | 0.074 |
| **V9K** | 8.878 | 0.061 |
| **V10K** | 8.83 | 0.035 |

### 3.6 Steric effects of amino acid substitutions

The structural alignment analysis showed that variants V9K and V10K produced the largest local RMSDs of 8.878 Å and 8.830 Å, respectively. On the other hand, the hydrophobic amino acid substitutions produced the lowest RMSDs of 1.452 Å and 1.580 Å for V8L and V10L, respectively positively correlating to the morphing results obtained. Analyzing the global RMSD of the NTD, due to these substitutions, the largest global RMSD was recorded by V8K (RMSD; 0.074 Å), followed by V9L (RMSD; 0.062 Å) and V9K (RMSD; 0.061 Å). The rest of the variants recorded less than 0.046 Å (Table 3). This suggests that at the global level of the NTD of the protein, the largest structural perturbation caused due to the amino acid changes occurred at the amino acid position 8 when Valine is changed to lysine.

In contrast, at the local regions, amino acid position 9 produced the most significant conformational alteration when Valine was changed to lysine. In a buried hydrophobic side chain, for which a comparatively smaller side-chain as in Valine, was replaced by a larger one as in leucine or isoleucine, we expect steric effects; however, we expect less change in this hydrophobic region as compared to the introduction of a charged amino acid such as lysine or aspartic acid. Thus, we observed, as in Table 3, that there were comparably smaller local and global RMSDs recorded in the hydrophobic amino acid variants (V8I, V8L, V8L). This could suggest that substituting a hydrophobic moiety into a buried hydrophobic region might not substantially alter the stability of the protein or might even contribute to the stabilization of the protein. Similar results have been recorded by Dong et al., 2008 **(25)**.

### 3.7 Quantitative analysis of variant TSPs electrophoresed under non-denaturing condition

To investigate the contribution of the hydrophobic valine patch on the structural stability of the NTD of P22 TSP, the hydrophobic valine patch was mutagenized to substitute either charged amino acid with Valine or hydrophobic amino acid with Valine. The created mutants were Val-to-Ile, Val-to-Leu, Val-to-Asp, Val-to-Lys using site-directed mutagenesis (**10**), and these mutants were expressed, and the tailspike proteins electrophoresed under native conditions. As shown in Figures 2A and 2B, both hydrophobic amino acids substituted mutants (Val-to-Ile, Val-to-Leu) showed no significant difference between them and WT native species. Our previous communication demonstrated that they also migrated at the same molecular level in native conditions **(10)**. Under native conditions, Val-to-Lys substitution at the three different positions (V8K, V9K, and V10K) did not produce any differences between these variant species and the WT native species (Figure 2C). Substituting Valine for the negatively charged aspartic acid (V8D and V10D) showed interesting results (Figure 2D). Both substitutions produced a higher amount of the intermediate species than the native species of the protein, and these were significantly higher amount of the intermediate species in these two mutants than all the variants produced under these native conditions. The migration of V8K and V10K at the molecular size of the intermediate might point to denatured NTD of the P22 TSP; such a finding has been proven **(10)**.

**Figure 2.**
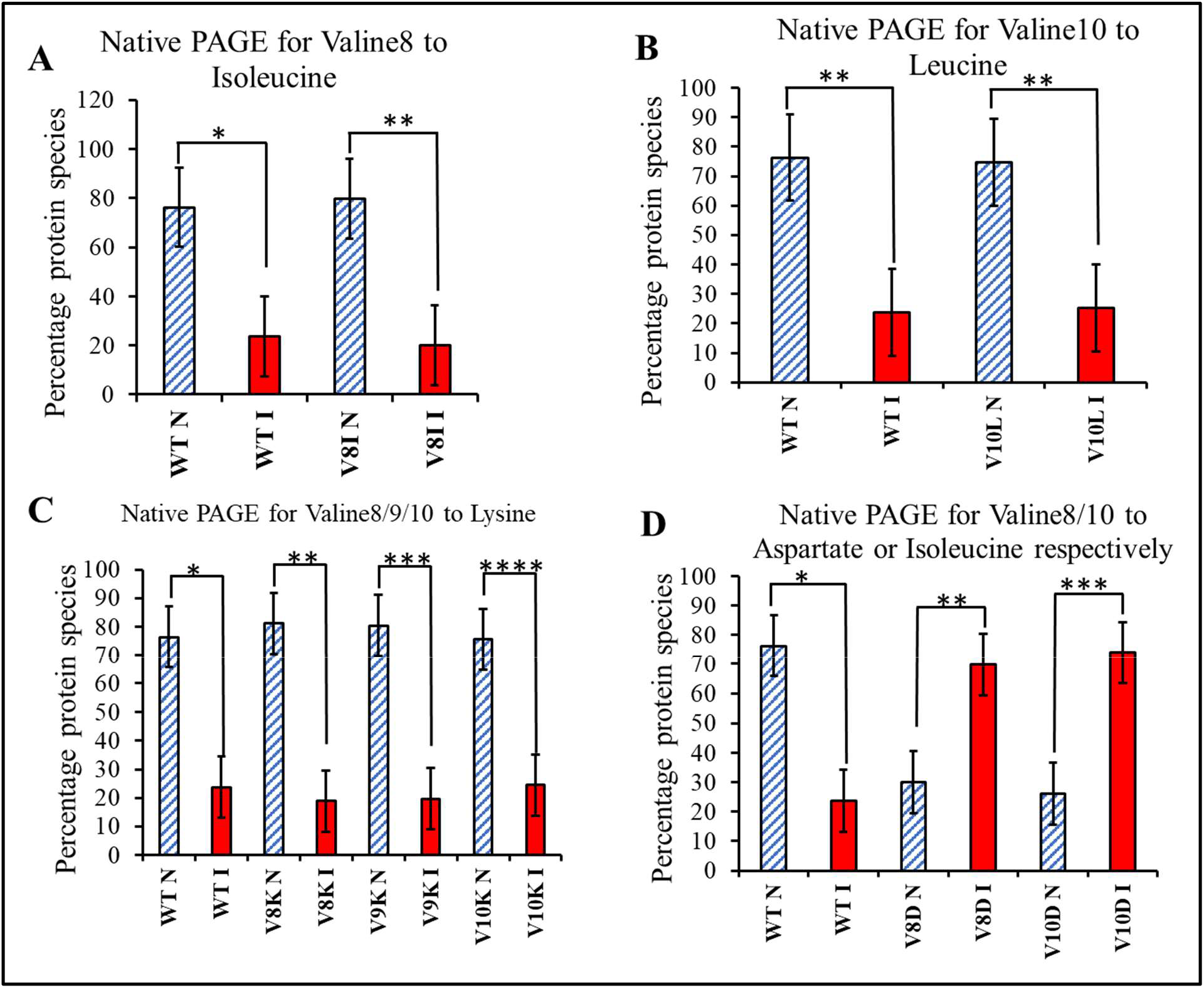
Quantitative analysis of variant TSPs electrophoresed under non-denaturing condition (SDS-free conditions, via native-PAGE). WT N = Wildtype P22 TSP native species. WT I = Wildtype P22 TSP intermediate species. V8I N= Valine-to-Isoleucine substitution variant TSP native species. V8I I= Valine-to-Isoleucine substitution variant TSP intermediate species. V10L N = Valine-to-Leucine substitution variant TSP native species. V10L I= Valine-to-Leucine substitution variant TSP intermediate species. V8K = Valine-to-Lysine substitution variant TSP. V8D = Valine-to-aspartic acid variant TSP. n = 3; *, **, ***, and **** p-value < 0.05.

### 3.8 Quantitative analysis of variant TSPs electrophoresed under denaturing condition

Although the trajectories of both the native and intermediates of the variant TSPs were demonstrated under the non-denaturing conditions, such understanding underscores the properties of these proteins under denaturing conditions such as detergent or heat. To investigate the contribution of the valine patch to the structural stability of the variant TSPs under denaturing conditions, the mutant proteins were mixed with SDS 2% final concentration and incubated at room temperature. Then proteins were electrophoresed using an electrophoretic buffer containing 2% SDS in an SDS-PAGE. The migration of these proteins under SDS denaturation has been demonstrated in our earlier publication **(10)** thus, providing the preamble for this quantitative analysis.

As shown in Figures 3A and 3B, under SDS denaturation conditions at room temperature, the protein species of both hydrophobic substitute variants (V8I and V10L) showed a significantly high amount of native state conformation species similar to the wildtype native species. This was expected since, hydrophobic-to-hydrophobic substitution might not significantly affect the hydrophobicity of the region and possibly maintain a hydrophobic core as the wildtype.

**Figure 3.**
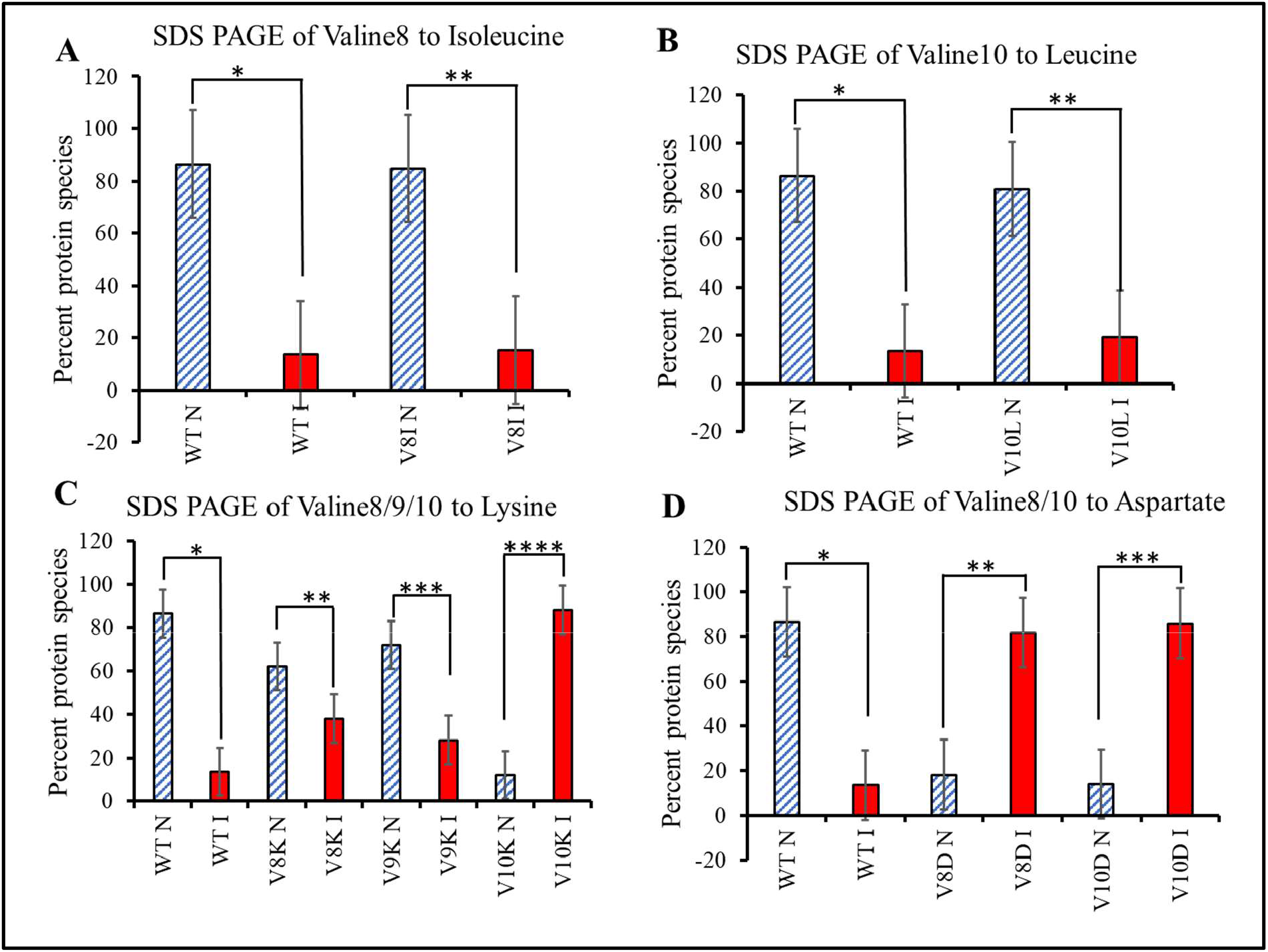
Quantitative analysis of variant TSPs electrophoresed under denaturing condition (2% SDS, via SDS-PAGE). WT N = Wildtype P22 TSP native species. WT I = Wildtype P22 TSP intermediate species. V8I N= Valine-to-Isoleucine substitution variant TSP native species. V8I I= Valine-to-Isoleucine substitution variant TSP intermediate species. V10L N = Valine-to-Leucine substitution variant TSP native species. V10L I= Valine-to-Leucine substitution variant TSP intermediate species. V8K = Valine-to-Lysine substitution variant TSP. V8D = Valine-to-aspartic acid variant TSP. n = 3; *, **, ***, and **** p-value < 0.05.

However, in the Valine to charge residue substitution as in V8K, V9K, V10K, V8D, and V10D, as shown in Figure 3C and 3D, there were significant differences between the wildtype species and the variants. The variant V8K recorded an average species abundance of 61.98% for the native conformers as compared to 38.02% abundance for the intermediate species under SDS (Figure 3C). V9K native species recorded 71.82%, whereas the intermediate recorded 28.18% (Figure 3C). There was a reversal of species abundance in the V10K under SDS denaturation; the native species (11.93%) of V10K fell significantly lower than the intermediate species (88.07%). Thus indicating that Valine-to-Lysine substitution at position 10 was the most structurally disruptive mutation of the study.

For V8D variants, the native species recorded a low of 18.25%, whereas 81.75% were recorded for the intermediate species. Similar trends were observed for the V10D variant, with a 14.11% native species abundance compared to 85.89% intermediate species abundance. Such findings demonstrate that substituting Valine to an aspartic acid negatively impacts the stability of the protein against denaturants such as SDS. Further work is necessary to validate and assess the extent to which the NTDs’ property is compromised due to these mutations; however, it is certain that these mutations led to the denaturation of the NTD of the trimeric protein **(10, 26)**.

The tailspike protein of the *Salmonella* phage P22 has been shown to be highly resistant to both SDS and heat **(19, 27, 28, 29, 30)**. However, in a combination of both denaturing conditions, the protein can be denatured at lower temperatures **(10)**. P22 TSP in the presence of SDS (2%) at 65 °C-75 °C has been shown to form an SDS-thermal intermediate (I), which subsequently disassociates into the individual monomers **(10, 19)**. This study carried out an *in vitro* thermal-SDS denaturation kinetics at 70 °C, 2% SDS. As shown in Figure 4A, V8D showed a highly unstable variant protein, with the majority of the protein species falling to the intermediate conformation state. A similar trend is observed with V10D variant (Figure 5B). The hydrophobic substitution variants (V8I (Figure 4B) and V10L (Figure 5C)) showed higher concentrations of the native species at the start of the incubation at 70 °C, 2% SDS, however, with increasing time of incubation at this condition, the native conformations dropped continually to form more intermediate species. Thus higher abundance of intermediate species is observed with increasing incubation time. For instance, while at 0-time point V8I, registered 80.41% native species, after 120 minutes of incubation in 70 °C, 2% SDS, the native species fell to 53. 75%. Sharper fall of native species was observed in the V10L variants at 120 minutes (registering native species abundance of only 45.51%) (Figure 5C).

**Figure 4.**
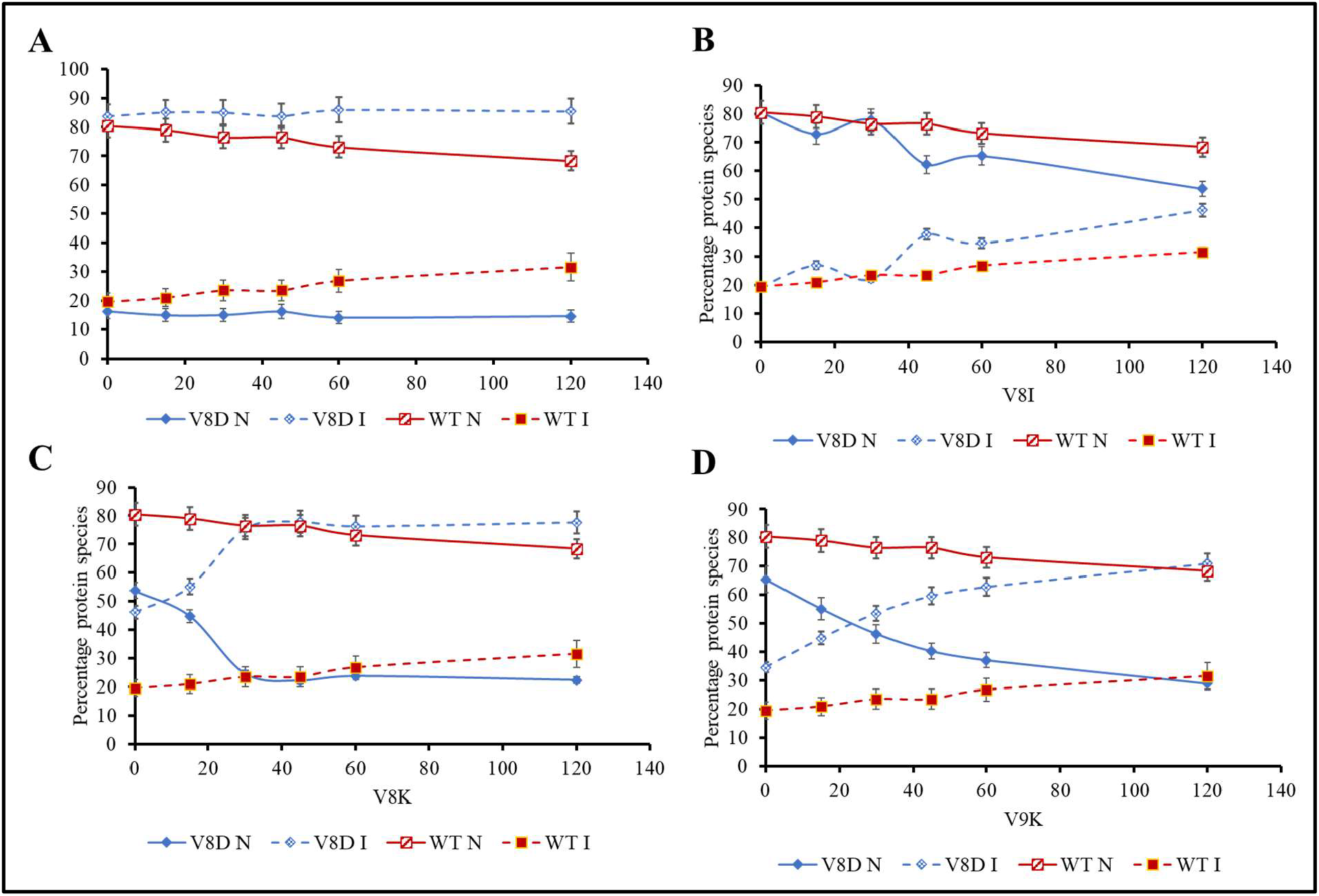
Kinetic analysis of variant TSPs electrophoresed under thermal-SDS denaturing condition (2% SDS, 70 °C). WT N = Wildtype P22 TSP native species. WT I = Wildtype P22 TSP intermediate species. V8I N= Valine-to-Isoleucine substitution variant TSP native species. V8I I= Valine-to-Isoleucine substitution variant TSP intermediate species. V8K = Valine-to-Lysine substitution variant TSP at amino acid position 8. V10K = Valine-to-Lysine variant TSP, change made at the amino acid position 9. The number of replicates was 3. Samples were taken at the set time points for 120 minutes.

**Figure 5.**
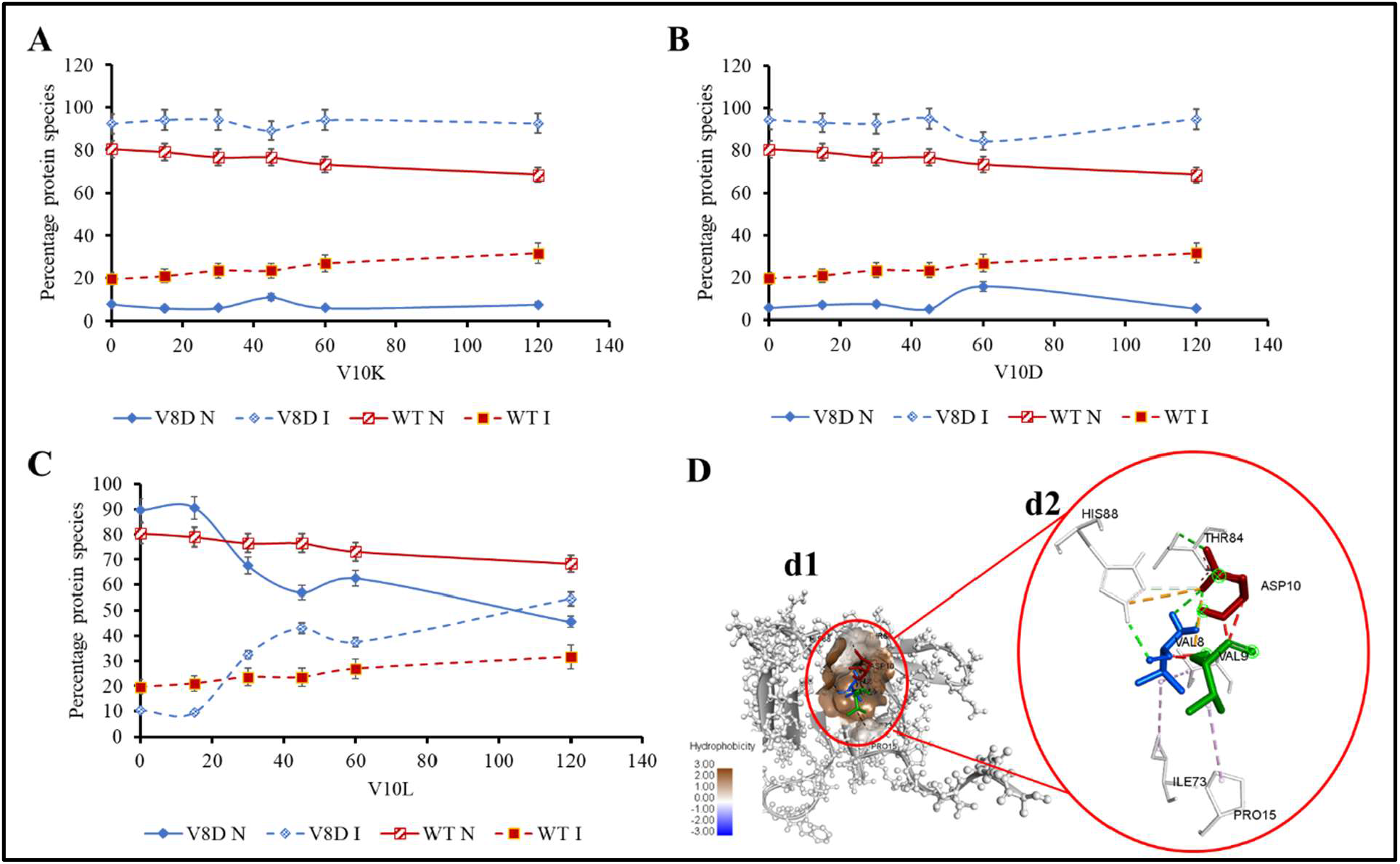
Kinetic analysis of variant TSPs electrophoresed under thermal-SDS denaturing condition (2% SDS, 70 °C). **(A)** WT N = Wildtype P22 TSP native species. WT I = Wildtype P22 TSP intermediate species. V10K N= Valine-to-Lysine substitution variant TSP, native species. V10K I= Valine-to-Lysine substitution variant TSP, intermediate species. **(B)** V10D = Valine-to-aspartic acid substitution variant TSP, at amino acid position 10. **(C)** V9L = Valine-to-Leucine variant TSP, change made at the amino acid position 10. Number of replicates = 3. Samples were taken at specific time points for 120 minutes. (D) Specific interaction of the mutated site (V10D) with the interacting chain in the hydrophobic core of the NTD of the V10D variant protein. (V10D, d1) shows the VVD epitope of the V10D variant docked to the hydrophobic region of the interacting chain (level of hydrophobicity indicated in brown color). (Figure 5D, d2) Shows the details of the interaction between the three amino acid epitope (VVD) and the adjacent interacting chain.

For Valine-to-Lysine mutants, (V8K (Figure 4C) and V9K (Figure 4D)), the both registered an initial native species abundance of 53.78% and 65.41%, respectively at the 0-minute time point; however, these species continued to fall in abundance until there was 22.35% and 28.99% after 120 minutes of incubation under this condition. The gradual and continued accumulation of intermediates indicates that 70 °C-2% SDS could provide adequate unfolding conditions to denature these variants into their intermediate forms. However, at this condition, after 120 minutes, there were no monomeric species formed. V10K (Figure 5A) showed that species were at the intermediate state at the start of the experiment and indicated no dissociation into monomers even after 120 minutes of heating at 70 °C in 2% SDS. It is interesting to observe that although these species had their NTD denatured, and yet they were still resistant to SDS and heat at 70 °C since they continued to persist as trimeric intermediates.

In Figure 5D, we showed the specific interaction of the mutated site (VVD of the V10D variant) with the interacting chain in the hydrophobic core of the NTD of the V10D variant protein, showing that the valine-valine-aspartic acid (VVD) epitope of the V10D variant docked to the hydrophobic region of the interacting chain (Figure 5D, d1). The level of hydrophobicity indicated that epitopes were fully buried in a hydrophobic core. As shown in Figure 5D, d2, the details of the interaction between the three-amino acid epitope (VVD) of the variant (V10D) and the adjacent interacting chain in the NTD of the protein demonstrates that this epitope interacted with the adjacent chain via electrostatic and hydrogen bonds (See Table 4).

**Table 4.** The details of the interaction between the three-amino acid epitope (VVD) of the V10D variant and the adjacent interacting chain of the NTD of P22 TSP, showing the bond distances and the type of interactions occurring.

| Residue | Interactions | Bond category | Bond distance/ Å |
| --- | --- | --- | --- |
| D10 | HIS88 - ASP10 | Electrostatic | 4.50 |
| D10 | THR84- ASP10 | Hydrogen Bond | 2.18 |
| D10 | THR84 - ASP10 | Hydrogen Bond | 2.62 |
| V8 | HIS88 - VAL8 | Hydrogen Bond | 2.31 |
| D10 | HIS88 - ASP10 | Hydrogen Bond | 3.36 |
| **V9** | PRO15 - VAL9 | Hydrophobic, Alkyl | 4.68 |
| **V8** | VAL8 - ILE73 | Hydrophobic, Alkyl | 5.30 |
| **V8** | VAL8 - ILE82 | Hydrophobic, Alkyl | 4.56 |

### 3.9 Comparing the variant models with WT NTD of P22 TSP

Models of the various mutant structures were subjected to structural alignment with the WT NTD of P22 to reveal subtle structural differences. As shown in Figure 6, apart from the valine patch region (in which amino acid changes were made), there were no observed major structural differences between the variants and the WT proteins, which is not surprising since most of the global RMSDs recorded for all the variants showed a very minimal difference from the WT (See Table 3).

**Figure 6.**
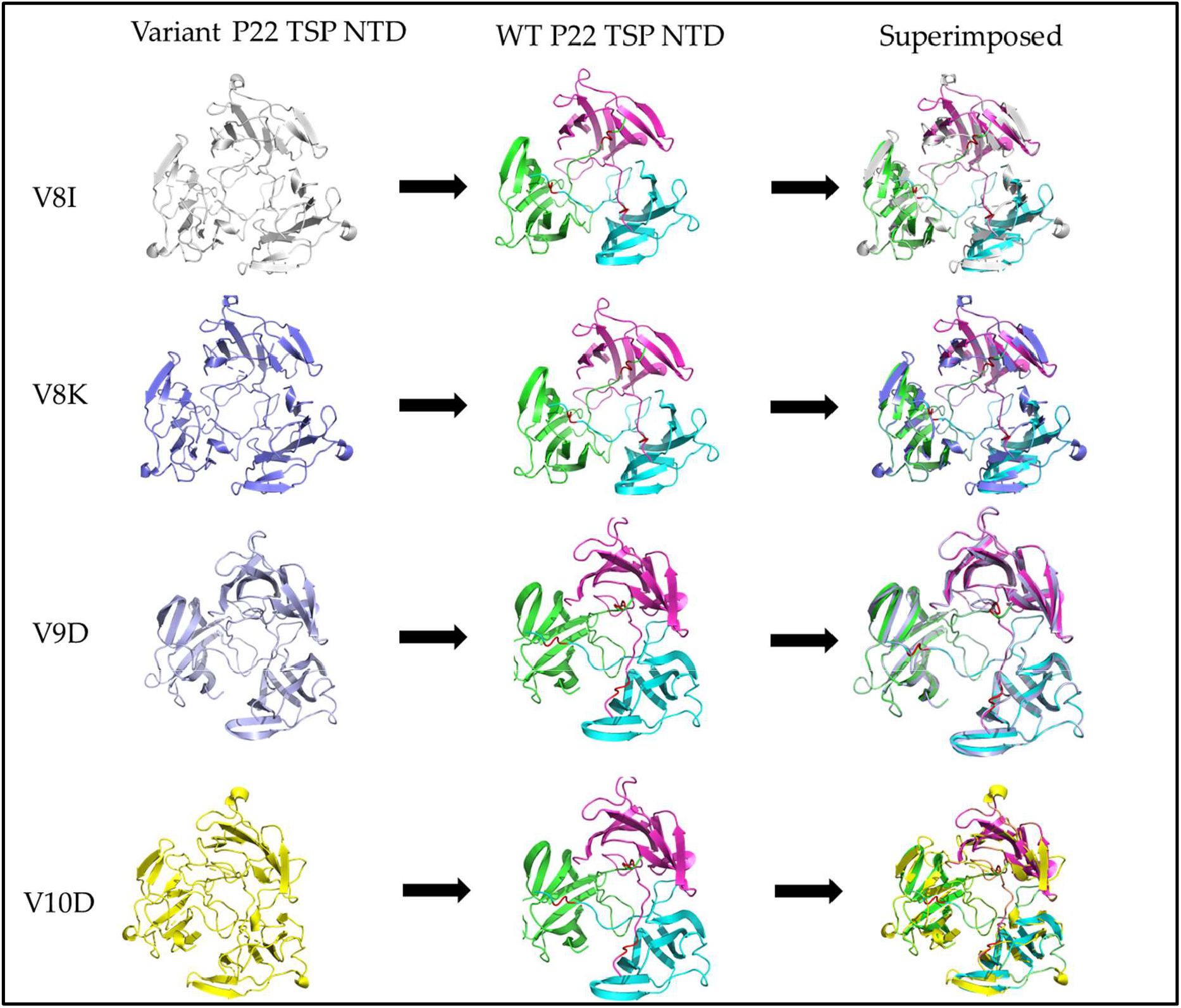
Alignment of variant models to WT P22 TSP NTD. Structural alignment created using PyMol 2 (Schrödinger, INC. Wild-type NTD chain A (Green), Chain B (Pink), Chain C (Blue). Site of site directed mutagenesis has been colored red.

### 3.10 Thermal denaturation of variant TSPs at 90 °C

As shown in Figure 7, the thermal denaturation of TSP-WT (Upper panel) and TSP-V8I (lower panel) at 90 °C only demonstrated that the WT TSP possesses higher thermal stability than the V8I variant TSP. As shown, heating the WT-TSP at 90 °C produced intermediates at the 20-minute time point, which continued to accumulate with time to produce approximately 50% intermediates at the 1-hour time point. The V8I variant TSP showed a high accumulation of intermediates at a 10-minute time point, and after 30 minutes, the intermediates, as well as the native species were completely disappeared (dissociated into monomers). As shown in Figure 8, the thermal denaturation of V8K TSP at 90 °C only demonstrated that the V8K variant TSP completely denatured within 10 minutes; thus, no native trimeric species nor intermediates are observed after 10 minutes.

**Figure 7.**
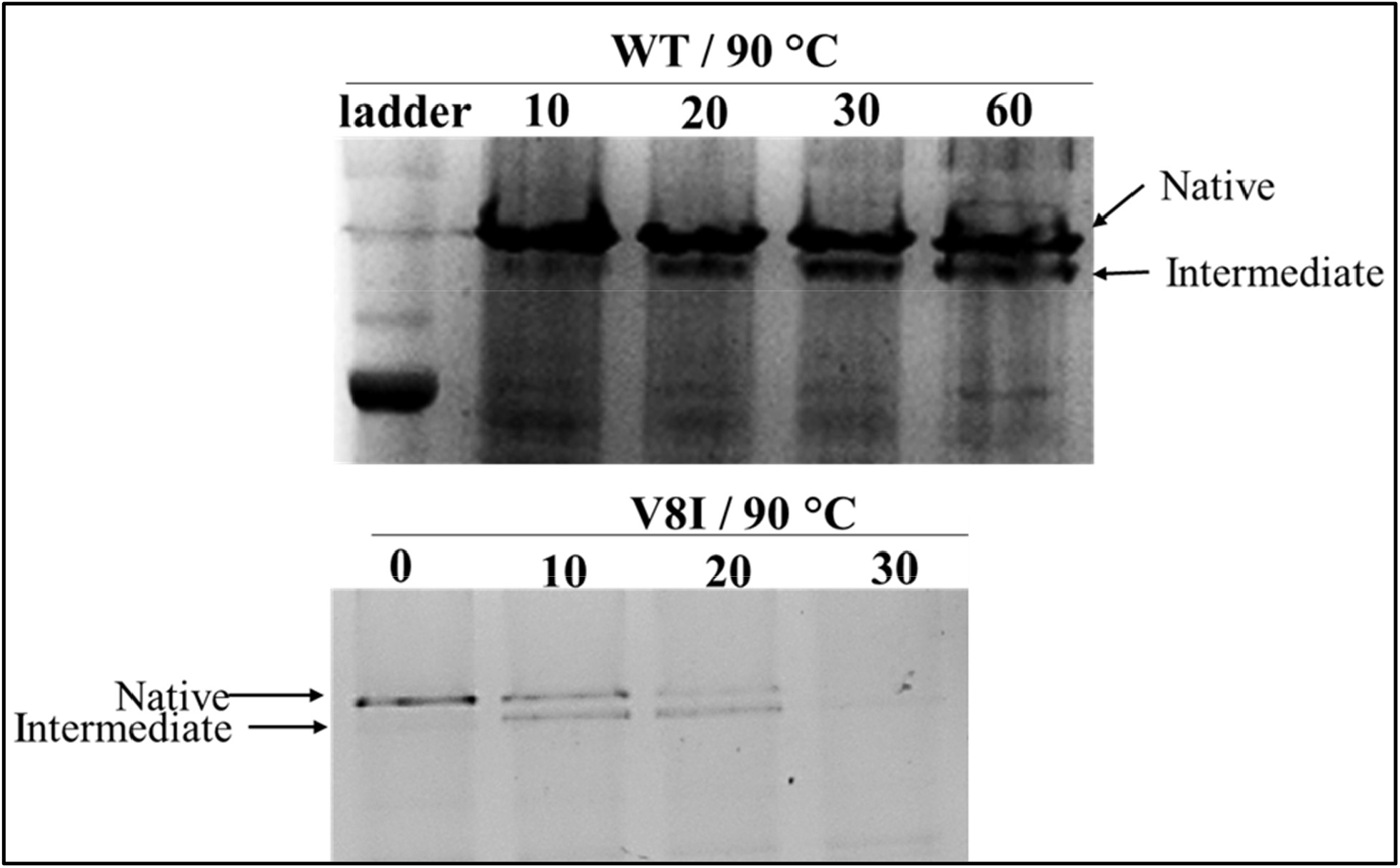
Thermal denaturation of TSP-WT (Upper panel) and TSP-V8I (lower panel) at 90 °C only. As shown, heating the WT-TSP at 90 °C produced intermediates at the 20-minute mark which continued to accumulate with time to produce approximately 50% intermediates at the 1-hour time point. The V8I variant TSP, showed a high accumulation of intermediates at 10-minute time point and after 30 minutes the intermediates as well as the native species had completely disappeared (dissociated into monomers).

**Figure 8.**
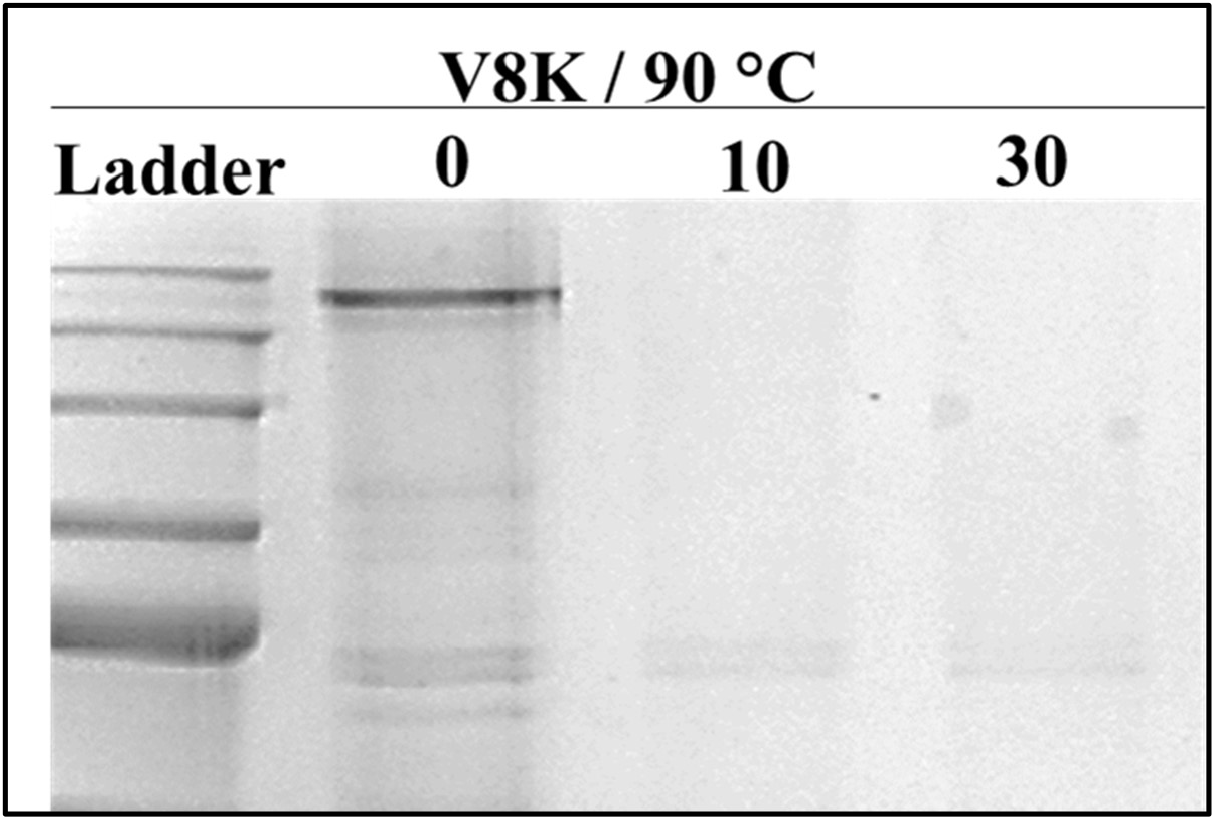
Thermal denaturation of V8K TSP at 90 °C only. As shown, heating the V8K variant TSP completely denatured the species within 10 minutes, thus no native trimeric species nor intermediates are observed after 10 minutes.

### 3.11 Thermal denaturation of variant TSPs at 70 °C

As shown in Figure 9, the thermal denaturation of TSP-WT (Upper panel) and TSP-V8I (lower panel) at 70 °C only demonstrated that heating the WT-TSP at 70 °C did not produce any intermediates after an hour. In contrast, the V8K variant produced observable intermediate species band within the first 10 minutes of heating at 70 °C, and after 30 minutes, all native species of V8K had been chased into the intermediate species. This indicates that the V8K variant is less thermal-stable than the WT TSP.

**Figure 9.**
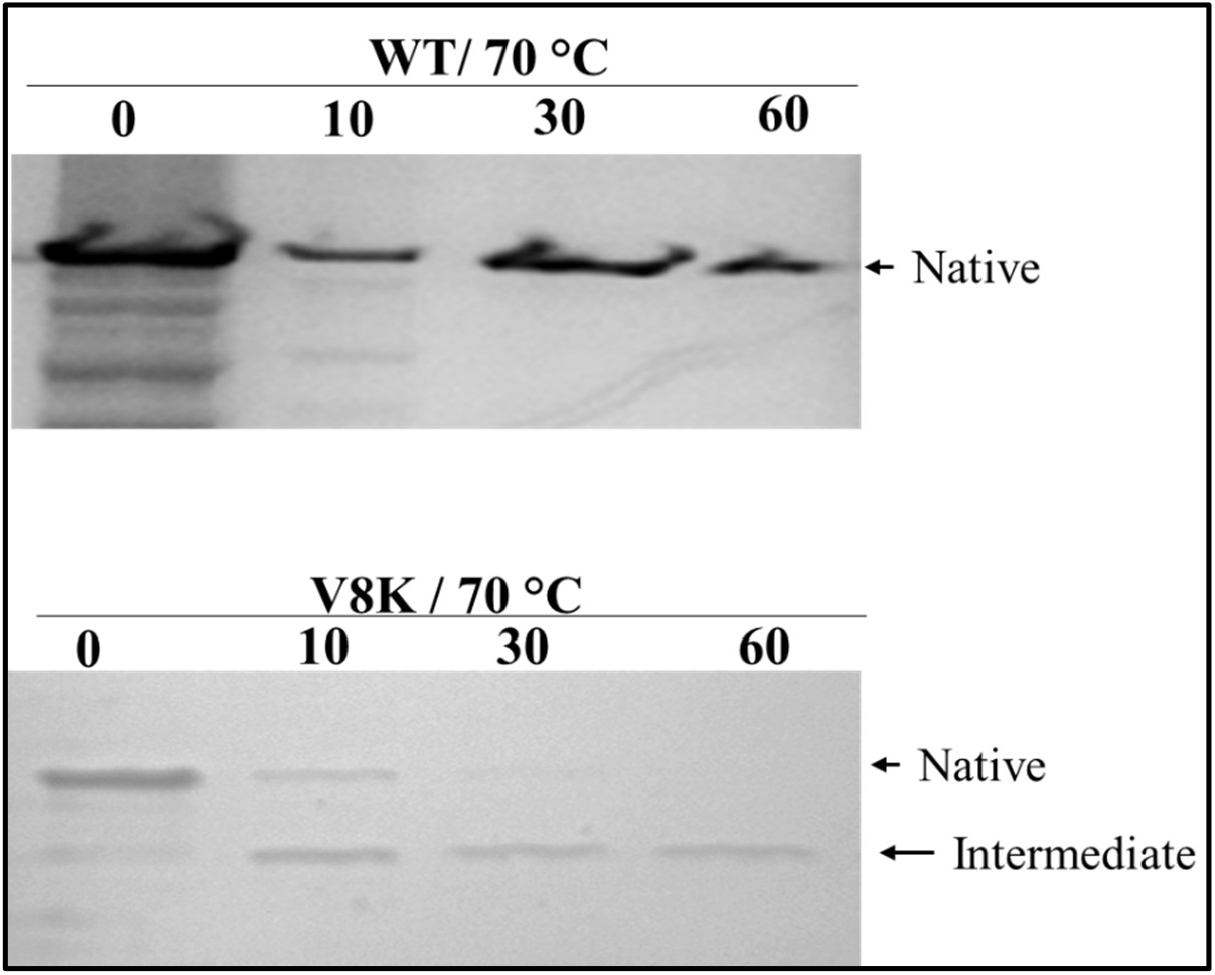
Thermal denaturation of TSP-WT (Upper panel) and TSP-V8I (lower panel) at 70 °C only. As shown, heating the WT-TSP at 70 °C did not produced any intermediates after the 60 minutes of heating, whereas V8K variant produced an observable intermediate species within the first 10 minutes of heating at 70 °C, and after 30 minutes all native species of V8K had been chased into the intermediate species.

## Conclusions

Several published works have demonstrated that hydrogen bonds and hydrophobic patches play crucial roles in the folding and stability of proteins **(31, 32, 33)**.

In this work, we confirmed the validity of this knowledge by computational and *in vitro* analysis of variant NTD of P22 TSP. We first showed that substituting Valine with aspartic acid and lysine produced the most dramatic changes in global motions of the TSP. This also correlated with lower stability of the variant TSPs to denaturation conditions such as SDS and heat. Comparatively, aspartic acid substitution produced the denatured NTD of the P22 TSP, and this has been confirmed by others **(10, 11, 19)**. Substituting Valine in the valine patch with hydrophobic residues such as leucine or isoleucine demonstrated comparably higher thermostability and SDS resistance compared to the charged amino acid substituted variants. These observations indicate that hydrophobic substitution into hydrophobic regions did not significantly disrupt the wildtype structural configurations of the TSP and the protein’s dynamics at room temperatures. However, compared to the WT TSP, both hydrophobic mutants showed lower thermostability, nonetheless far greater than the charged amino acid substituted mutants, thus supporting the notion that hydrophobic to hydrophobic substitution in proteins exhibit similar overall characteristics. Thus, this work confirms our previous conclusion that the trivalent hydrophobic patch (V8-V9-V10), located on the stem peptide, is a crucial player in the stability of the entire P22 NTD. This crucial knowledge is essential in understanding the molecular interactions occurring during P22 phage assembly during the life cycle of the phage and thus might be vital for bioengineering purposes in which P22 TSP can be manipulated to achieve tunable affinity for the rest of the phage or chimeric protein that can be used for biosensing, pathogen immobilization, or streptavidin-coated magnetic beads for ELISA.

## Acknowledgements

We wish to thank past members of the Villafane’s laboratory for their contributions and input into this work. We are grateful to Alabama State University and the College of Science, Mathematics and Technology for their continuous support.

