## Supplementary Figure S1 to S10 for "Single Amino Acid Change Mutation in the Hydrophobic Core of the N-terminal Domain of P22 TSP affects the Proteins Stability": Supplementary information for Single Mutation paper.docx

**Figure S1**. Close view of the conformational dynamics of V8D NTD. Shown in red is the mutagenized valine patch.

**Figure S2**. Conformational dynamics of the entire V8D NTD. Shown in red is the mutagenized valine patch. Chain A shown in green, Chain B in pink and Chain C in blue.

**Figure S3**. Conformational dynamics of the entire V9D NTD. Shown in red is the mutagenized valine patch. Chain A shown in green, Chain B in pink and Chain C in blue.

**Figure S4**. Conformational dynamics of the entire V10D NTD.

**Figure S5**. Close view of the conformational dynamics of V10D NTD.

**Figure S6**. Conformational dynamics of the entire V8K NTD.

**Figure S7**. Conformational dynamics of the entire V9K NTD.

**Figure S8**. Conformational dynamics of the entire V10K NTD.

**Figure S9**. Conformational dynamics of the entire V8I NTD.

**Figure S10**. Conformational dynamics of the entire V8L NTD.
