## Supplementary figures and images for "Single Amino Acid Change Mutation in the Hydrophobic Core of the N-terminal Domain of P22 TSP affects the Proteins Stability"

### Figure S1. V8D Upclose.gif

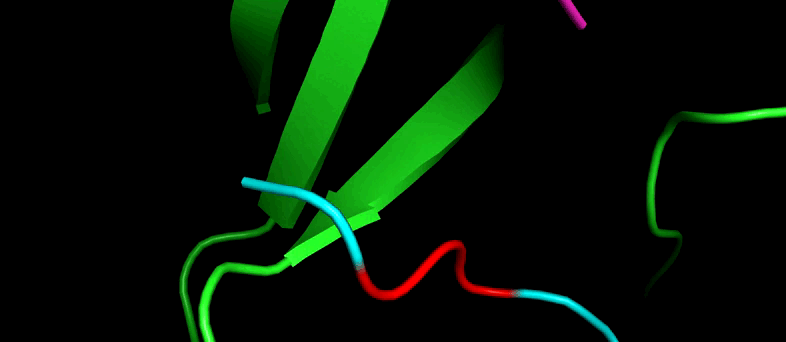

### Figure S2. V8D.gif

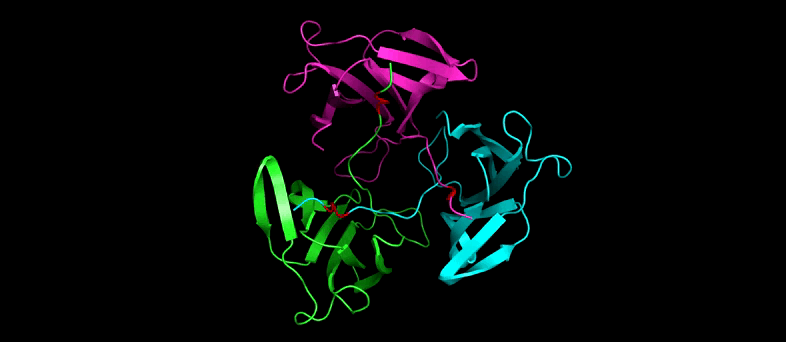

### Figure S3. V9D.gif

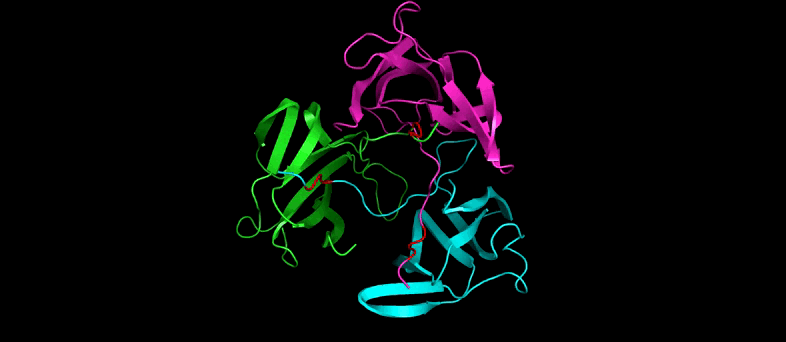

### Figure S4. V10D again.gif

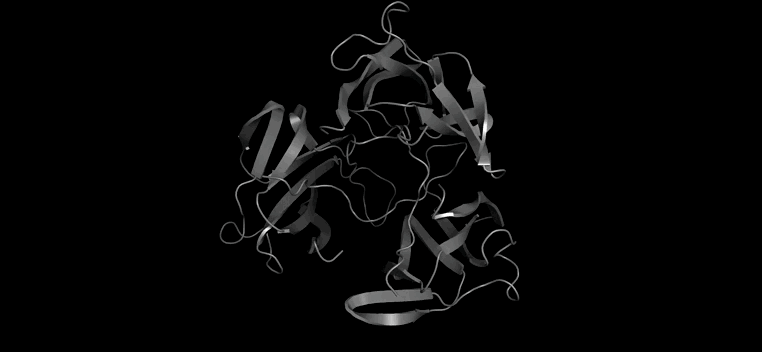

### Figure S5. V10D upclose.gif

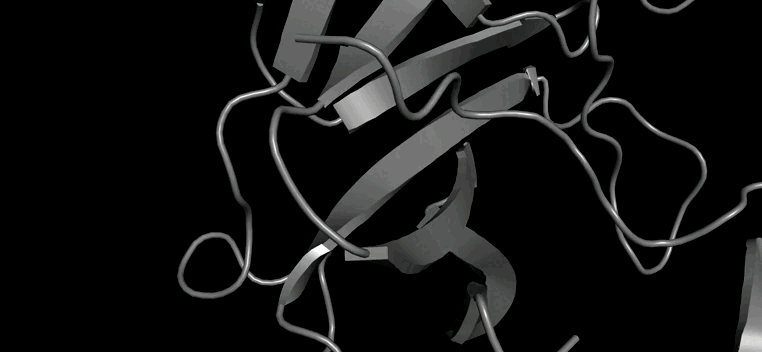

### Figure S6. V8K Colored.gif

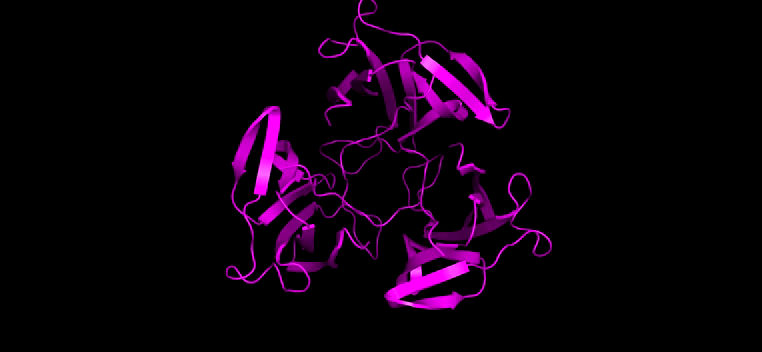

### Figure S7. V9K Colored.gif

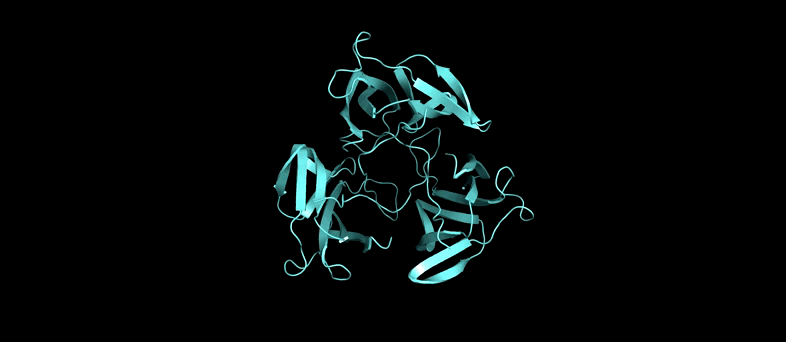

### Figure S8. V10K.gif

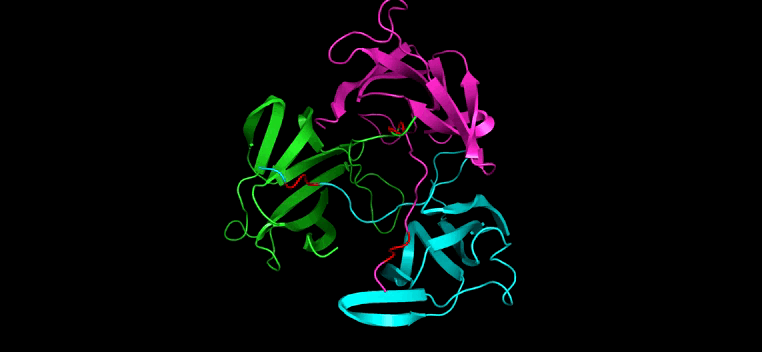

### Figure S9. V8I orange color.gif

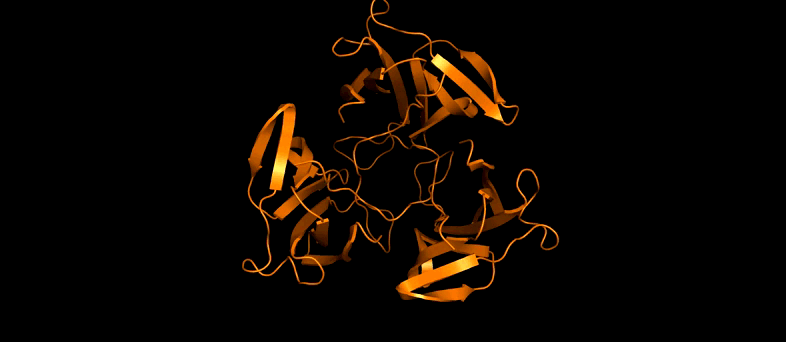

### Figure S10. V8L colored brown.gif

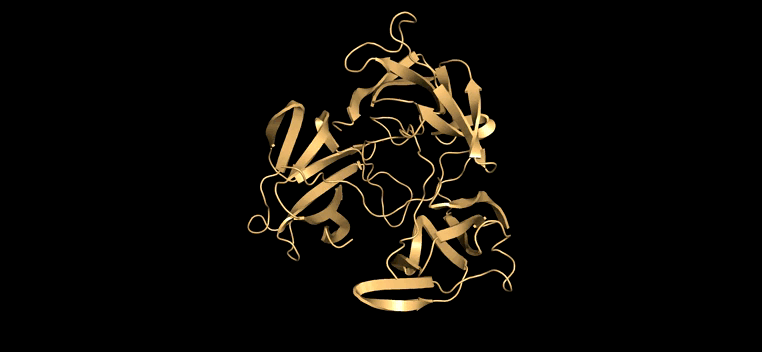
